# Versatile Tissue-Injectable Hydrogels with Extended Hydrolytic Release of Bioactive Protein Therapeutics

**DOI:** 10.1101/2023.09.01.554391

**Authors:** Eric S. Nealy, Steven J. Reed, Steve M. Adelmund, Barry A. Badeau, Jared A. Shadish, Emily J. Girard, Fiona J. Pakiam, Andrew J. Mhyre, Jason P. Price, Surojit Sarkar, Vandana Kalia, Cole A. DeForest, James M. Olson

## Abstract

Hydrogels generally have broad utilization in healthcare due to their tunable structures, high water content, and inherent biocompatibility. FDA-approved applications of hydrogels include spinal cord regeneration, skin fillers, and local therapeutic delivery. Drawbacks exist in the clinical hydrogel space, largely pertaining to inconsistent therapeutic exposure, short-lived release windows, and difficulties inserting the polymer into tissue. In this study, we engineered injectable, biocompatible hydrogels that function as a local protein therapeutic depot with a high degree of user-customizability. We showcase a PEG-based hydrogel functionalized with bioorthogonal strain-promoted azide-alkyne cycloaddition (SPAAC) handles for its polymerization and functionalization with a variety of payloads. Small-molecule and protein cargos, including chemokines and antibodies, were site-specifically modified with hydrolysable “azidoesters” of varying hydrophobicity via direct chemical conjugation or sortase-mediated transpeptidation. These hydrolysable esters afforded extended release of payloads linked to our hydrogels beyond diffusion; with timescales spanning days to months dependent on ester hydrophobicity. Injected hydrogels polymerize *in situ* and remain in tissue over extended periods of time. Hydrogel-delivered protein payloads elicit biological activity after being modified with SPAAC-compatible linkers, as demonstrated by the successful recruitment of murine T-cells to a mouse melanoma model by hydrolytically released murine CXCL10. These results highlight a highly versatile, customizable hydrogel-based delivery system for local delivery of protein therapeutics with payload release profiles appropriate for a variety of clinical needs.

**Translational Impact:** We developed injectable hydrogels that provide loco-regional, controlled release of protein therapeutics. Local delivery is especially suitable for potent, protein-based drugs like chemokines and monoclonal antibodies whose systemic toxicities can be life threatening. Our hydrogel is equipped with slowly hydrolyzing linkers for transient payload coupling, prolonging its therapeutic window and minimizing the need for repeat surgeries in a clinical setting. A range of release profiles, spanning days to over a month, and a broad compatibility to therapeutically relevant, recombinant proteins provide clinicians with flexible therapeutic options to suit a variety of circumstances.

## 1. Introduction

Delivery of therapeutic agents directly into tissue or body compartments allows clinicians to achieve higher local drug concentrations than could be achieved by systemic administration, which is hindered by dose-limiting toxicities (DLTs).^1–5^ Hematologic, hepatic, and neurologic effects are common examples of DLTs that may prevent achieving effective drug concentrations following systemic delivery.^6,7^ This has led to the development of novel local or regional delivery strategies aimed at minimizing DLTs and improving therapeutic efficacy. Loco-regional delivery may be particularly suited for protein-based therapeutics, such as monoclonal antibodies and cytokines. Although they have high potency and target specificity, their systemic administration can cause damage to healthy tissues, particularly due to on-target/off-cancer or similar toxicities.^8–17^

Hydrogels, a class of biomaterials comprised of water-swollen polymer networks, are commonly used to deliver therapeutics locally to the tissue surrounding their implant site. These materials can swell with fluid, facilitating the exchange of nutrients and molecules between tissue and the gel.^18–32^ Synthetic hydrogels, like those made of poly(ethylene glycol) (PEG), are minimally immunogenic and can be polymerized and functionalized with therapeutic payloads using bioorthogonal click chemistry.^18,19,23–35^ These gels offer unique flexibility, with mechanical properties and environmental responsiveness that can be adjusted to suit various biological settings. However, there are drawbacks, particularly in terms of the consistency and longevity of therapeutics delivered. Many polymer-housed therapeutics rely on simple diffusion or structural breakdown to release payloads into tissue, leading to short-lived therapeutic exposure and uneven drug release at the shrinking implant-tissue interface.^4,5,20,36,37^ Addressing these concerns is a necessity in clinical settings, especially in sensitive organs like the brain where repeat surgeries are typically not an option.

Our team has developed a tunable hydrogel platform to overcome the limitations of inconsistent therapeutic exposure and short-lived payload release. Using bioorthogonal SPAAC click chemistry, our system enables customizable release rates and broad compatibility with therapeutically relevant proteins without compromising their bioactivity.^38–40^ We genetically encoded recombinant protein therapeutics, like chemokines and antibodies, with C-terminal sortase-recognition sequences (i.e., LPXTG, where X is any amino acid), which allowed for site-specific modification with azide-functionalized polyglycine peptides (PolyG-azidoester) via chemoenzymatic transpeptidation.^25,41,42^ Recombinant CXCL10-LPETG modified with PolyG-azidoesters were linked to a hydrogel and injected once into a murine melanoma model. This resulted in similar T-cell recruitment and suppressed tumor growth compared to multiple injections of soluble chemokines purchased from a vendor. Our results suggest the use of PolyG-azidoester linkers with differing hydrolysis rates enables the customization of local protein therapeutic release from hydrogels in a “fire-and-forget” manner, with a variety of user-defined release rates achievable to suit different clinical needs.

## 2. Results

### Characterization of PEG-tetraBCN Hydrogels with User-Defined Payload Release

Our study aims to showcase the utilization of four-arm PEG-tetraBCN hydrogels (M_n_ ≈ 20,000 Da) as an exceptionally customizable and bioorthogonal platform for protein therapeutic delivery.^24,27^ The hydrogel network is formed through polymerization via strain-promoted azide-alkyne cycloaddition (SPAAC), employing a PEG-diazide crosslinker (M_n_ ≈ 3,400 Da) (**Figure 1A**). SPAAC is a click reaction that proceeds optimally at 37°C in aqueous solutions and exhibits high compatibility with various chemical moieties and bioactive molecules, including commercially available azidoacids.^24,27^ These SPAAC-compatible fatty acids were utilized in our studies as linkers between the mock fluorescent payload DEAC-OH (MW = 247.29 Da), and the PEG-tBCN backbone (**Figure 1B**).^43^ Ester bonds formed between the azidoacid and the payload hydrolyze at rates dependent on the length of the azidoacid’s carbon chain, matching trends with what has been reported with similar linker chemistries. ^44^ Therefore, the hydrolysis rate of a therapeutic payload from these gels can be customized to meet clinical needs by selecting preferred azidoacid lengths (**Figures 1C-D**). We demonstrated this customizability *in vitro* by quantifying the release of DEAC-OH from a hydrogel coupled with 2-azidoacetic, 3-azidobutanic, or 4-azidopropanic acids to yield “DEAC-2,3,4-azidoesters”. We analyzed the release of the payload, indicated by supernatant fluorescence, from 10µL gels immersed in phosphate-buffered saline (PBS) at 37°C via spectroscopy over 4 weeks. Our results showed that ∼80% of unlinked DEAC was released within 24 hours, meanwhile ∼60% of the DEAC-4azidoester payload was released by day 23 (**Figure 1C**). Release differed according to the azidoacid used, potentially bestowing user-defined control over this hydrolytic release system. We showcase this by simultaneously esterifying DEAC with 2,3, or 4- azidoacids, then quantifying its release into PBS from 10µL gels. Our results show complete DEAC release achieved within 5 days using 2- azidoacetic acid, 21 days for the 3-azidobutanic acid, and only ∼69% released using the 4-azidopropanic acid by day 25 (**Figure 1D**).

**Figure 1.**
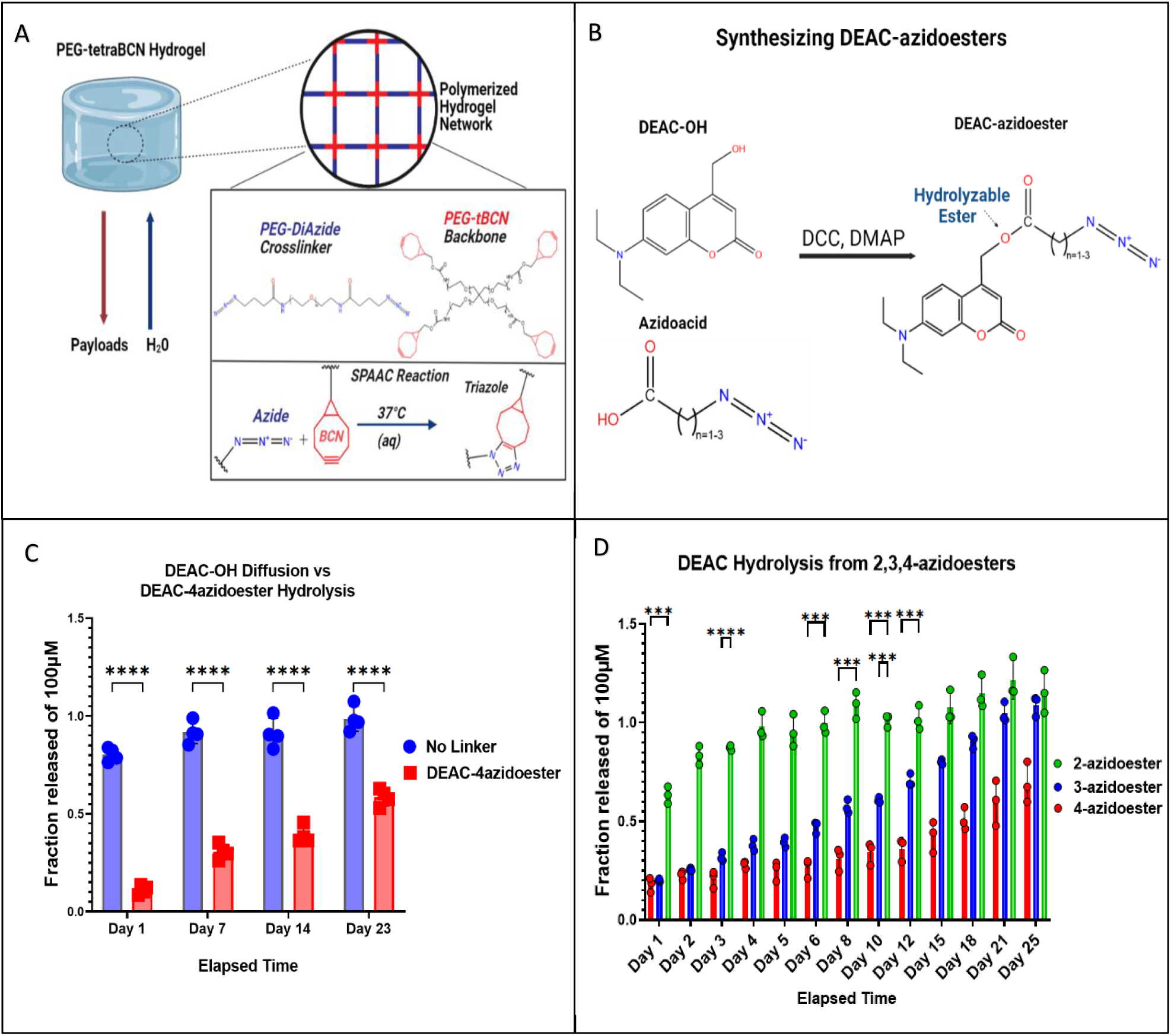
Characteristics of a Bioorthogonal Hydrogel with Tunable Payload Release Rates. A) Illustration depicting the components of a PEG-tetraBCN hydrogel and its individual reactants. The 4-armed PEG-tetraBCN backbone is labeled red and the linear PEG-diazide crosslinker is in blue. The temperature sensitive SPAAC reaction between a BCN group and an azide result in a covalent triazole adduct, polymerizing the gel. B) Steglich esterification converts DEAC-OH and 2,3,4-azidoacids into DEAC-azidoester.^1^ Ester bonds linking DEAC to an azidoacid hydrolyze in aqueous environments. C) Comparison of DEAC-OH diffusion release profile from a hydrogel versus DEAC-4azidoester hydrolysis from a hydrogel. Data are presented as mean ± SD for n=4 replicates. Statistical significance was determined by multiple unpaired t-tests followed by Holm-Šídák posthoc correction. D) Release assay demonstrates the tunability of azidoester system, with 2-azidoesters completely releasing rapidly and 4- azidoesters releasing over multiple weeks. Statistical significance was determined by two-way ANOVA followed by followed by Holm-Šídák posthoc correction. Only displaying p-values < 0.001. [(***) p < 0.001, (****) p < 0.0001]

### Adapting Azidoester Linkers for Proteins via Functionalization onto GGGGRS Peptide Adapters

To achieve programmed and sustained release of a protein therapeutic using an azidoacid linker, a chemical modification involving an ester bond is required. Therapeutically relevant proteins, from small cytokines to large monoclonal antibodies, may have numerous sites that are reactive with esterification chemistry **(Figures 2A-B)**. Chemical modification of the protein with an azidoacid on its own is not appropriate, as random attachments on a functional protein can lead to disrupted biological activity or uncleavable amide tethering to the gel.^23,25^ Therefore, the “H-GGGGRS-NH_2_” polypeptide was synthesized, whose C-terminal serine can be esterified with an azidoacid to create “PolyG-azidoester” **(Figures 2C-D)**.^43^ The N-terminal GGG-can then be enzymatically attached to any recombinant protein encoded with terminal LPXTG motif via sortase-mediated coupling (“sortagging”).^41,45^ Pre-esterifying serine (or threonine / tyrosine)-containing polypeptides, like H-GGGGRS-NH_2_, with an azidoacid of a desired length/hydrolysis rate, can enable the rapid generation of a suite of adapter molecules that allow the transient linkage of a protein therapeutic to the backbone of a SPAAC hydrogel.

**Figure 2.**
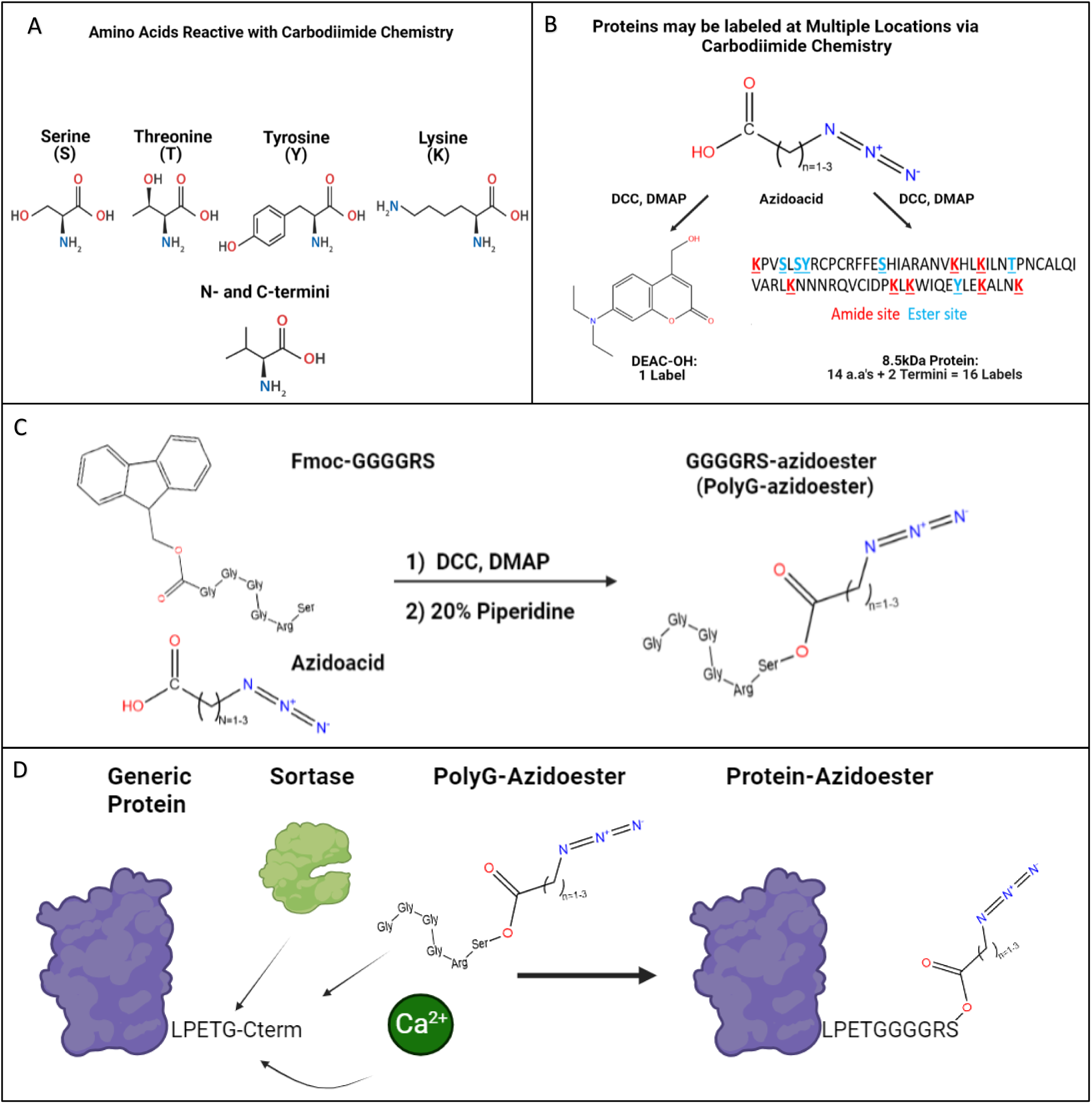
Synthesizing Polypeptide Adapter Molecules to Facilitate Protein Incorporation into SPAAC Hydrogels. A) Illustration showing all possible ester and amide chemical conjugation sites on amino acids and N/C-termini. B) When chemically conjugating azidoacids to DEAC-OH, there is only one location that is reactive. On the other hand, protein payloads typically have multiple coupling sites that may result in uncleavable amide linkage or disrupted protein function. C) Site-specific azidoester linkage on proteins can be achieved by using a polypeptide adapter molecule pre-functionalized with an azide group and chemoenzymatically appended onto the N/C-termini of a recombinant protein containing an LPXTG sortase recognition site. D) Schematic demonstrating the “sortagging” of a PolyG-azidoester to a generic protein to create a Protein-azidoester capable of binding to a SPAAC hydrogel.

### Expressing and Sortase Tagging Protein-Based Immunomodulators with SPAAC-Compatible GGGGRS-Azidoesters

One significant advantage of utilizing the sortase tagging technique to bestow SPAAC hydrogel compatibility lies in its broad applicability to various recombinant proteins and expression systems. In this context, we showcase two distinct methods for expressing recombinant, sortase-taggable proteins in mammalian and bacterial systems: Traditional sortase tagging and Sortase-Tagged Expressed Protein Ligation (STEPL).^41,42^ What we refer to as “Traditional” sortase tagging follows the original method described by Popp et al. This method was employed with a mammalian cell-expressed IgG modified with sortase motifs on its C-termini (IgG-LPETG 2x) **(Figure 3A)**. ^41^ Sortase 5M enzyme, separately expressed in E.coli, was employed to modify the IgG-LPETG (2x) with two PolyG-azidoesters **(Figure 3B)**. Western blot confirms the removal of the 6xHis tag from IgG-LPETG Sortagged with PolyG-4azidoester and its ability to flow through a Ni-NTA column. Unmodified or partially modified IgG-LPETG were obtained from Ni-NTA elution and maintain reactivity with the 6xHis antibody **(Figure 3C)**. Furthermore, we can confirm the presence of an azide conjugated to the IgG in this process, and that the 6xHis tag was not simply cleaved off at the LPETG motif without transpeptidation, as sortase is capable of under certain circumstances.^47,48^ We conducted an overnight click reaction between both IgG-LPETG and IgG- 4azidoester with DBCO-IRDye 680. DBCO is another molecule with ring strain that is compatible with SPAAC click chemistry ^49^. Western blot analysis revealed a 680nm signal, approximately the expected size of an IgG (∼150kDa), exclusively in the lane containing IgG-4azidoester **(Figure 3D)**.

**Figure 3.**
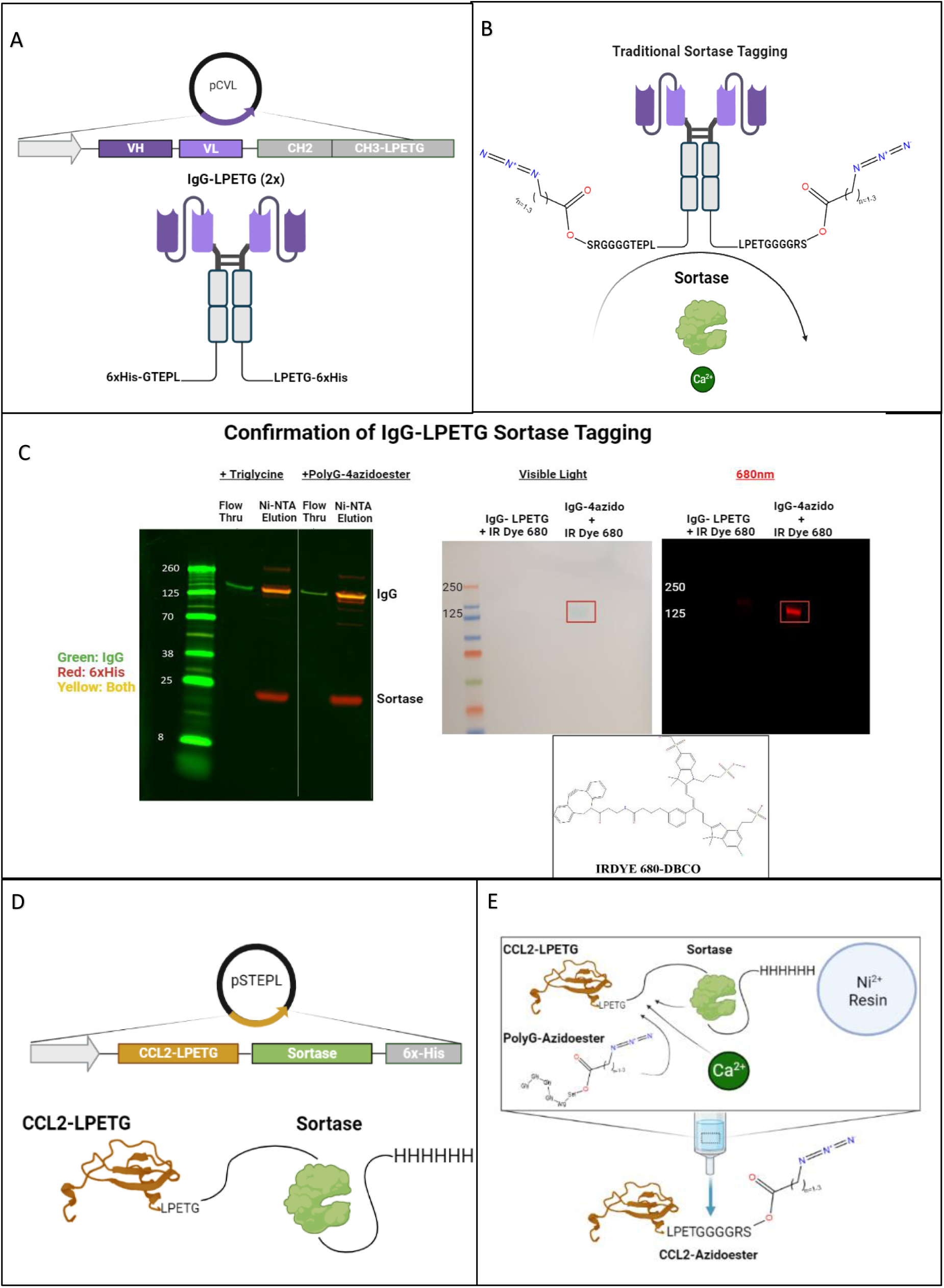
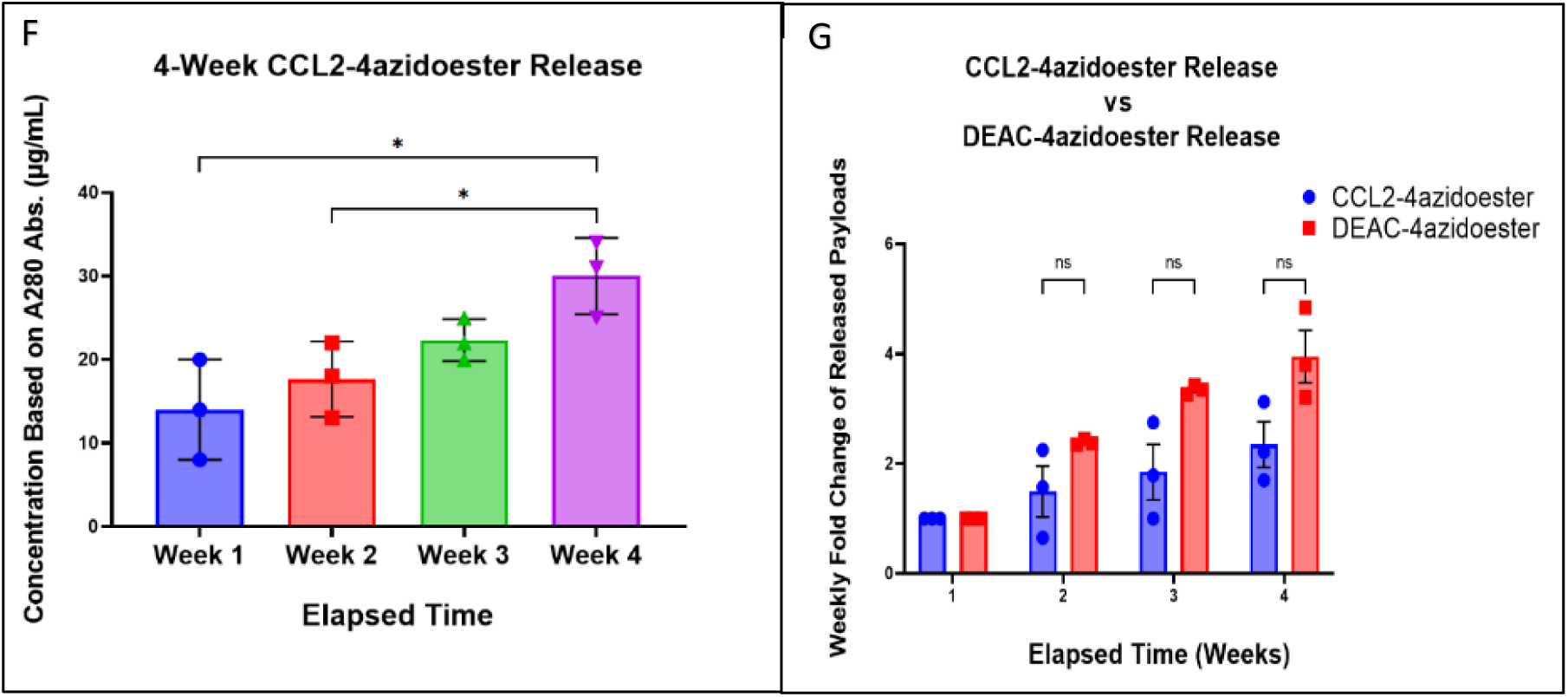
Expression and Sortagging of Immunomodulatory Proteins with Hydrolysable Azidoesters. A-B) Illustrations of IgG-LPETG expression via the Daedalus mammalian expression platform and its modification with PolyG-4azidoesters.^2^ “Traditional” sortase tagging was utilized, requiring separate expression of sortase and the protein target of interest prior to tagging with GGG-containing species.^3^ C) The Western blot confirms the removal of both 6xHis tags (red channel) from IgG-LPETG (green channel) after sortagging with a triglycine positive control or PolyG-4azidoester. Sortagged proteins flow through a Ni-NTA column, while unmodified or partially modified proteins with 6xHis tags remain attached to the column. (Right) Another Western blot compares the parental IgG-LPETG with IgG-4azidoester after overnight reaction with DBCO-IRDye 680. An accumulation of dye appears at the MW of an IgG (∼150kDa) in the IgG-4azidoester lane only. Scanning this blot in the 680 (red) channel confirms a signal from the dye at the MW of IgG-4azidoester. D) This illustration depicts the structure of a CCL2-STEPL fusion protein expressed via the STEPL system in E. coli.^4^ In this variant of sortagging, the protein of interest and sortase are co-expressed on the same plasmid. E) This illustration represents the STEPL process as follows: Purified fusion proteins are initially bound to a Ni-NTA column. Upon adding PolyG-azidoester and calcium, sortase catalyzes the simultaneous removal of CCL2 and the attachment of PolyG-4Azidoester, resulting in CCL2-4Azidoester. This product can be collected in the flowthrough, while the remaining fusion protein continues to be bound to the Ni-NTA column until elution. F) This graph displays month-long release profiles of CCL2-4-azidoester from a hydrogel in PBS at 37°C, quantified as µg/mL by A280 readings. Data are presented as mean ± SD for n=3 replicates. Statistical significance was determined by one-way ANOVA followed by Tukey’s multiple comparison test. G) In this graph, fold change of CCL2-4-azidoester release over 4 weeks is compared to that of DEAC-4azidoester. Despite CCL2’s much larger size, we observed no significant difference in release rates between the two species. Data are presented as mean ± SE for n=3 replicates. Statistical significance was determined by multiple unpaired t-tests followed by Holm-Šídák posthoc correction. [(ns) not significant, (*) p < 0.05.

The azidoester hydrolysis-mediated release system offers a means to establish and maintain a concentration gradient over a desired timeframe, which is essential for the function of some secreted proteins, like chemokines. Chemokines have emerged as promising adjuncts to immunotherapy, particularly when dealing with immunosuppressed tumors. These molecules play a pivotal role in guiding immune cell trafficking to sites of infection and tissue infarction.^50–53^ We chose C-C motif chemokine ligand 2 (CCL2), a chemoattractant for monocytes and other immune cells, as a representative therapeutic protein for expression within the STEPL system.^54–60^ We then utilized this protein as a model for sustained protein release *in vitro* after conjugation with PolyG-4azidoester. CCL2 was first expressed as a STEPL fusion protein in E. coli, enabling the simultaneous expression of our chemokine of interest and sortase on the same plasmid. **(Figure 3D, S1)**. In contrast to IgG-LPETG, which possesses two LPETG motifs, CCL2 contains a single C-terminal LPETG motif. Consequently, a 1:1 protein-to-linker ratio was expected to yield a release profile akin to DEAC-OH **(Figure 1C-D)**. The modified chemokine was released from the larger STEPL fusion protein through sortase in the presence of calcium and the PolyG-4azidoester linker, where it can be collected from the flow through **(Figure 3E, S2).** Hydrogel solutions containing CCL2- 4azidoesters were allowed to react overnight before undergoing polymerization, casting, and immersion into PBS at 37°C. Weekly A280 measurements of the released protein revealed a steady, albeit notable, increase in detected protein in the release medium over a 4-week period. In our hands, the fold change in CCL2-4azidoester release each week did not significantly differ from the trend observed with DEAC- 4azidoester in Figure 1 over four weeks, despite their significantly different sizes. This data suggests hydrolytic linker cleavage may be the rate-limiting factor for payload release of CCL2, but larger molecules may be more affected by diffusion. (**Figures 3F-G**).

### Hydrogel Solutions undergo *in situ* Polymerization within Living Tissue

We demonstrate the practicality of injecting pre-polymerized PEG-tetraBCN hydrogel solutions into living tissue without complex steps. By cooling the unpolymerized solution to 4°C, the temperature-sensitive SPAAC reaction is slowed, allowing sufficient time to aspirate and inject the pre-mixed solution using a silanized syringe (**Figures 4A-B)**.^61^ Clinicians may be able to inject amorphous hydrogel solutions into areas of the body where direct transplantation of a fixed 3-dimensional solid polymer is not feasible, such as the tight confines of the brain. We chilled a 6.5% (w/v) PEG-tetraBCN gel solution on ice, aspirated it into a Hamilton syringe, and successfully injected it into a living mouse brain (**Figure 4C**). After injection, the animal’s body temperature allows the SPAAC reaction to proceed at its optimal rate, and the newly polymerized gel forms within the needle tract of the injection site. The mouse survived for the entire 7-day study without any visible adverse effects from the gel in its brain, owing to inherent biocompatibility of the PEG gel components and SPAAC gelation chemistry.

**Figure 4.**
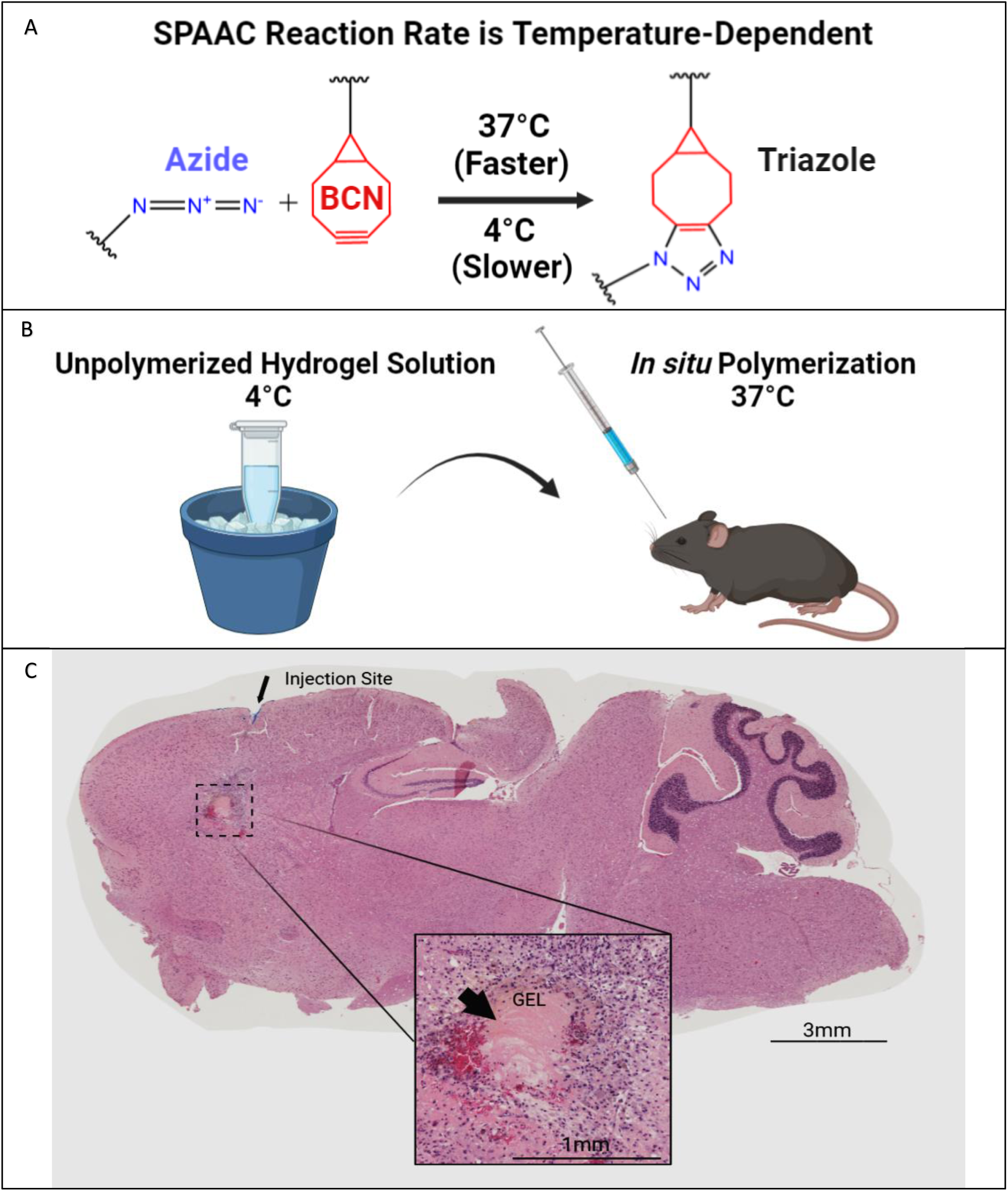
Injection and Polymerization of Hydrogel Solutions *in situ*. A) This illustration demonstrates the optimal temperature for the SPAAC reaction to proceed. B) Mixing the individual gel components on ice slows the SPAAC reaction, allowing additional time for aspirating the gel mixture into a syringe and prompt injection into an animal. The optimal polymerization temperature is reached upon exposure to physiological body temperature. C) A chilled PEG-tetraBCN hydrogel solution was administered into the frontal cortex of a living mouse using a silanized Hamilton syringe. After 7 days, the brain was harvested, sectioned, and later examined via H&E staining. The intact gel was found within the injection tract. Larger image taken at 10x magnification.

### Confirming Bioactivity of Hydrogel-released Murine CXCL10 in an *in vivo* Melanoma Model

Immunologically "cold" tumors often produce factors that can inhibit or disable T-cell infiltration. In this study, we demonstrate the translational potential of our hydrolysable release system by housing and delivering potent chemokine gradients into a "cold" mouse melanoma model to counteract T-cell exclusion. Specifically, we selected C-X-C motif chemokine ligand 10 (CXCL10), an interferon γ-induced T-cell chemokine, for this purpose. ^62–65^ We expressed murine CXCL10 in disulfide bond-capable E. coli and employed traditional sortase tagging to modify it with PolyG-3azidoesters. The 3-azidoacid was chosen for linkage because its expected release rate was best suited for 5 days of dosing. **(Figure S4)**. Upon completion of the study, we observed qualitative tumor growth attenuation and a significant increase in CD8+ T-cell trafficking in B16 melanoma flank tumors after injecting 4.5 µg of murine CXCL10-3azidoester tethered to hydrogels in a single dose **(Figures 5B-D, S5)**. To match the estimated protein release rate of 4-6% per day extrapolated from DEAC PolyG-3azidoester (**Figure 1D**), we administered 500ng doses of Peprotech murine CXCL10 solutions in PBS every other day over 5 days. Although this treatment group had a similar effect on CD8^+^ trafficking and tumor size, repeat administrations may not be practical in sensitive organs such as the brain, further supporting the use of a one-time hydrogel application. PBS and Hydrogel alone control groups showed no effect on T-cell recruitment or tumor volume, demonstrating that the preserved biological activity of the chemokines delivered by the hydrogel produced the observed results. Thus, we demonstrate a practical scenario in which locally delivering bioactive, protein-based therapeutics with user-defined release rates may be favorable.

**Figure 5.**
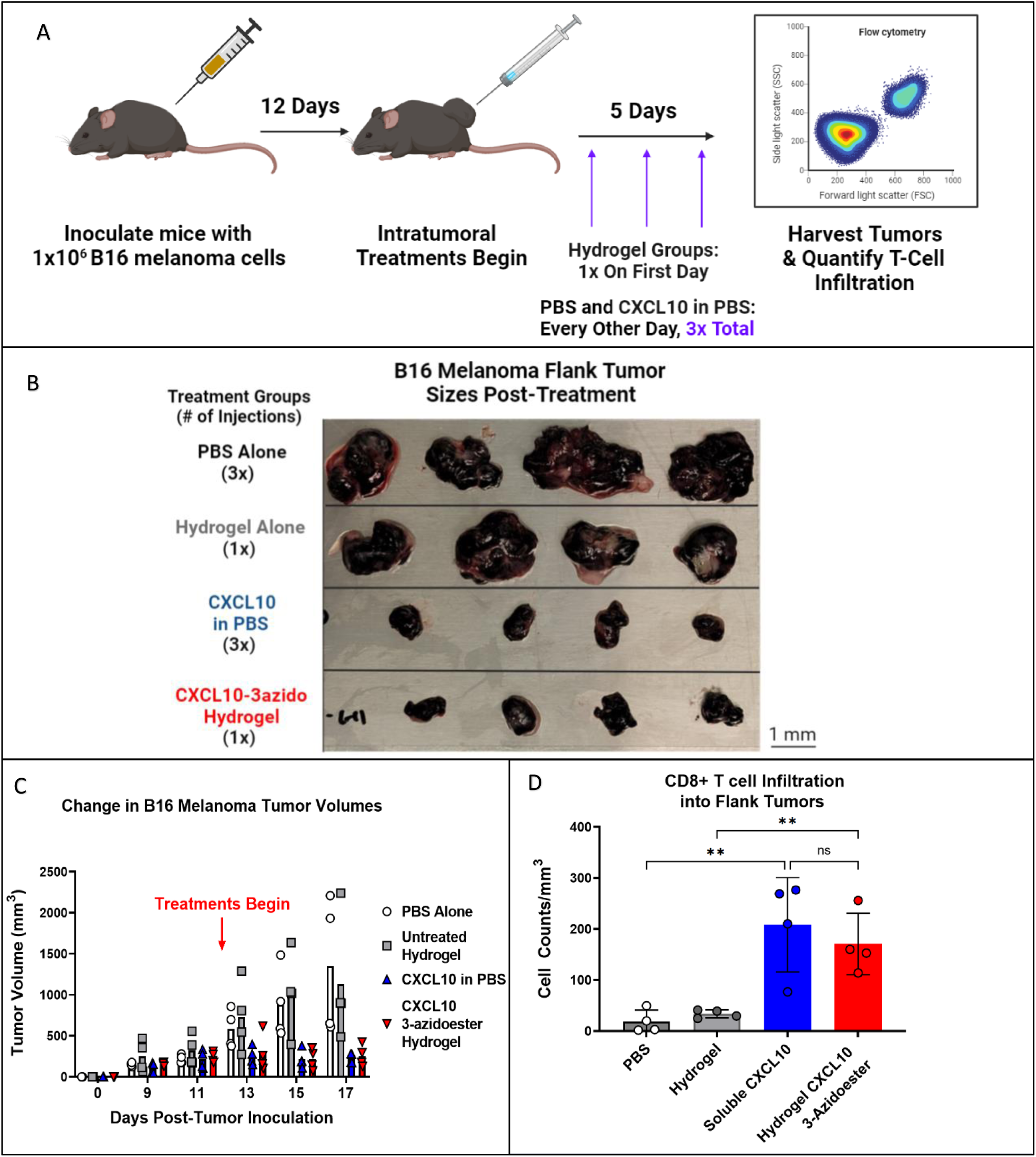
A Single Administration of a Hydrogel Housing CXCL10-3azidoester Recruits T cells into a “Cold” Tumor. A) This timeline illustrates the study design, including the timing of CD19^+^ B16 melanoma inoculations and the beginning of treatment administration: PBS Control, Hydrogel Control, Soluble CXCL10, and CXCL10- 3azidoester Hydrogel. B) This image shows the sizes of the flank tumors collected at the end of the study and demonstrates a stark contrast between treatment groups. C) CXCL10-treated tumors grew slower, while their matched controls grew faster (ns). Data are presented as mean ± SEM for n=4 replicates D) Quantification of CD8^+^ infiltrate into each tumor reveals a significant difference in CD8^+^ cells in tumors treated with Hydrogel CXCL10-3-azidoester compared to an empty hydrogel alone. Data are presented as mean ± SD for n=4 replicates. Statistical significance was determined by one-way ANOVA followed by Holm-Šídák posthoc correction. [(ns) not significant, (**) p < 0.01].

## 3. Discussion

Our study aims to develop highly customizable bioorthogonal PEG hydrogels that are both safe and effective in controlling the local delivery of bioactive protein therapeutics. To avoid the potential risk of systemic toxicity associated with soluble therapeutics, slow-release depots like hydrogels can provide sufficiently high concentrations of therapeutics within the tissue, while reducing the risk of systemic DLTs.^4,5,36,66,67^ We initially prioritized a high degree of biocompatibility in the construction of our material. We engineered a PEG-based hydrogel functionalized with SPAAC chemical handles, which is an enzyme-free variant of azide-alkyne click chemistry.^38^ PEG, a non-toxic and lowly immunogenic polymer, is currently employed in multiple FDA-approved applications, including therapeutics such as Neulasta and Movantik.^68,69^ SPAAC is a non-toxic click reaction that has demonstrated safety in the presence of living cells and enhances the promise of our hydrogel system for clinical applications.^30–32,70^ Furthermore, SPAAC reactions have shown promise in the construction of antibody drug conjugates utilized in clinical trials, including STRO-001 and ADCT-601.^71,72^

For the successful delivery of protein therapeutics via hydrogel, we firmly believe that extended-release capabilities are essential. This approach avoids the need for multiple surgeries and helps mitigate the toxicities associated with a burst release of drugs into tissue.^4,5,36,73^ One can achieve extended release simply by adjusting the gel’s crosslink density. The average pore size of the gel network is dependent on the length of the PEG chains and is directly related to diffusion rates out of the gel.^20,36,70,74^ However, it’s important to recognize that adjusting the diffusion rate of protein payloads, as demonstrated with insulin, may extend release rates on the scale of hours.^70^ This may not be sufficient for therapies meant to last many days or weeks. In contrast, the FDA-approved Oncogel polymer can sustain the release of paclitaxel over several months, similar to the release rate achieved with DEAC-OH on the 4-azidoester linker **(Figure 1C).**^36^ However, this prolonged diffusion rate primarily results from interactions between the hydrophobic drug and the co-block polymer, a situation not applicable to hydrophilic protein therapeutics within a hydrogel. Another strategy employed clinically is the slow physical degradation of the polymer to release payloads, as seen in the FDA-approved Gliadel wafer. Nonetheless, this approach can affect drug distribution profiles since the polymer doesn’t maintain the same contact with the tissue over time ^4,5,37,67,75^ And, despite its extended release capabilities, Oncogel, too, deteriorates over time. Ultimately, this results in similar challenges as those observed with Gliadel.

We believe our hydrolytic linker strategy is the superior option compared to those FDA approved methods for maintaining long-term, consistent protein release from an implanted depot. While the PEG-tBCN hydrogels used in our studies were not designed to lose structural integrity, the azidoacids utilized in the amidation of the PEG-diazide crosslinker could instead be functionalized as azidoesters. This would enable gel network hydrolysis over time at a predictable rate. There are longer azidoacids available commercially than those used in this study, potentially enabling protein release (or gel breakdown) timeframes far longer than what was observed in our experiments, if desired. Utilizing longer azidoacids in the hydrogel crosslinker may be useful in clinical situations where a foreign body response is a concern, but complete release of the gel’s contents is desired before depolymerization.

SPAAC polymerization and sortase-mediated conjugation strategies could be substituted for other click chemistry ligation strategies, protein modification techniques, and/or mechanisms of protein-linker cleavage. Click reactions, including oxime ligation, tetrazine ligation, nitrone dipole cycloaddition, and tetrazole photochemistry, can be substituted or used jointly with SPAAC.^76–82^ Hydrogel backbones and polypeptide adapters can be functionalized with these reactive handles based on preference. SpyTag/SpyCatcher, LOVTRAP, and HaloTag are among the site-specific protein modification techniques that can be used to substitute for “sortagging” to functionalize proteins to adapter molecules carrying click chemistry handles.^28,83–86^ We demonstrated with IgG-LPETG that two azides can be conjugated onto one protein, opening the door to heterogenous enzymatic tagging. This may be particularly applicable to “knob in holes” bispecifics, enabling linkage to the gel from one sortase motif and bioactive payload conjugation to another enzyme binding site. ^87^ Lastly, although our azidoester linker system provides a simple “fire-and-forget”, method of protein release, this mechanism can be more tightly controlled by clinicians by substituting hydrolytic azidoester linkages with light-responsive, drug-sensitive, Boolean-logic gate linkers.^24–27,83,88^

Based on our findings, we envision a scenario where heterogeneous payloads can be equipped with azidoesters of varying lengths. This approach would enable combinatorial therapeutic delivery with different release rates for each species, all from a single hydrogel depot. In summary, this versatile hydrogel-based, local delivery strategy has the potential to be a broadly adaptable and effective solution for a wide range of clinical requirements and scenarios.

## 4. Materials and Methods

### Synthesis and Validation of Hydrogel Components

#### PEG-tetraBCN Hydrogels

The individual hydrogel components used in these studies include: (1R,8S,9s)-bicyclo[6.1.0]non-4-yn-9-ylmethyl (2,5-dioxopyrrolidin-1-yl)carbonate (BCN-OSu), poly(ethylene glycol) tetrabicyclononyne (PEG-tetraBCN, M_n_ ∼ 20,000 Da), 2,5-dioxopyrrolidin1-yl 4-azidobutanoate (N_3_-OSu), and poly(ethylene glycol) diazide (PEG-diazide, Mn ∼ 3,400 Da). These components were synthesized and polymerized into PEG-tetraBCN hydrogels as previously reported in greater detail, including G’ and G”.^24,27,31^ In brief: 4-arm PEG-OH was functionalized with BCN-OSu (BCN-NHS ester) at a 1:4 molar ratio to create the PEG-tetraBCN gel backbone. Linear PEG-OH was functionalized with N_3_-OSu (N_3_-NHS ester) at a 1:2 ratio to create the hydrogel crosslinker, PEG-diazide. Hydrogels are polymerized by mixing PEG-tetraBCN backbones with PEG-diazide at 1:4 molar ratios at 37°C. Fully polymerized gels were formulated at concentrations of 8% (w/v) for *in vitro* studies and 6.5% (w/v) for *in vivo* injections as this concentration was easier to handle with the syringe.

#### DEAC 2,3,4-azidoesters and Release From PEG-tetraBCN Hydrogels

7-(Diethylamino)-4- (hydroxymethyl)coumarin (DEAC-OH) was previously synthesized in-house **(Method S3)** and conjugated to 2-azidoacetic acid (Click Chemistry tools, 1081), 3-azidopropionic acid (Synthonix, A1939), and 4-azidobutyric acid (Synthonix, A1941) via Steglich esterification. Briefly, 0.13 mmol of DEAC-OH, 0.30 mmol of DMAP, and 0.13 mmol of the corresponding azidoacid were mixed in minimal DCM and stirred for 10 minutes at room temperature. Next, 0.15 mmol of EDAC was added to the mixture, and the reaction was stirred overnight. The completed reaction was passed through a vacuum filter to remove urea byproducts and dried under vacuum. To prepare the hydrogel, 100 µM DEAC-azidoester conjugates were suspended in DMSO and “clicked” onto the backbone of PEG-tetraBCN hydrogel solutions overnight before polymerization. The resulting gels were cast as 10µL cylinders and plated in triplicate into a 12-well plate containing 500 µL PBS/well. The plate was incubated at 37°C and 5% CO_2_ for the duration of the experiment. At each timepoint, supernatants were collected from each well, and the fluorescence of the released DEAC-OH was measured using a Molecular Devices SpectraMax plate reader (Laser line 405nm; λ_ex_ 387nm, λ_em_ 470nm). The concentration of DEAC-OH was determined using linear regression of a DEAC-OH standard in PBS.

#### GGGGRS-3,4-azidoester (PolyG-azidoester)

The Fmoc-GGGGRS-NH_2_ polypeptide, which had C-terminal amidation, was either synthesized in-house **(Method S1-S2)**, or purchased from Biomatik Corporation (Ontario, CA). Fmoc-GGGGRS 3,4-azidoesters were produced using Steglich esterification, a type of carbodiimide chemistry. ^43^ Specifically, 0.45 mmol of Fmoc-GGGGRS, 2.0 mmol of DMAP (Sigma-Aldrich, 851055), and 0.74 mmol of 3 or 4 azidoacids were stirred for 10 minutes at 40°C in minimal dimethylformamide (Sigma-Aldrich, 319937). The solution was then added with 0.74 mmol of DCC (Sigma-Aldrich, D80002) and stirred overnight at 40°C. Following this, the polypeptide-azidoester was Fmoc-deprotected by adding piperidine (ChemImpex, 02351) to a final concentration of 20% and stirred for 5 minutes. Crude polypeptide-azidoester was then precipitated in cold di-ethyl ether, HPLC purified using 95:5 H_2_O/acetonitrile, and lyophilized. The completed products, with an appearance of a clear-yellow oil, was confirmed via MALDI-TOF mass spectrometry **(Figure S1)**. The purified “PolyG-azidoesters” were stored at -20°C under a nitrogen atmosphere for future use.

### Expression and Purification of Recombinant Proteins

#### IgG-LPETG (2x)

Though we do not present a binding or phagocytosis assays in this work, the VH/VH amino acid sequence of the 2.3D11 CD47mAb was obtained from US Patent US 9,650,441 B2. These sequences, along with CH2-CH3 of the IgG1 constant region were inserted into the pCVL-SFFV-muScn-IRES-GFP mammalian expression plasmid (Genscript, Nanjing, Jiangsu, China). Protein was expressed utilizing the Daedalus system with a C-terminal LPETG - 6x His tag.^46^ Proteins were purified and stored in PBS for later use.

#### CCL2-LPETG

The mature amino acid sequence for human CCL2 (aa 24-99) was obtained from NCBI, GeneID 6347. GBlocks (IDT) were created for this sequence with a C-terminal “LPETG” Sortase recognition site and complementary 5’ and 3’ overhangs to the NdeI/XhoI double digested pSTEPL Sortase fusion expression plasmid. ^42^ The resulting gBlocks were ligated into pSTEPL plasmids using Gibson assembly (NEB) and transformed into chemically competent SHuffle T7 Express E.coli (NEB). After bacterial liquid cultures were grown to 0.6 OD600, protein expression was induced overnight at 16°C with 0.2mM IPTG (Thermofisher, 15529019). The overnight cultures were lysed and sonicated in non-denaturing conditions: 20mM Tris, 125mM NaCl, 10mM imidazole, 0.1% Triton X-100 + COmplete tablet (Millipore Sigma, 11873580001), and the CCL2- LPETG STEPL fusion proteins were purified via Ni-NTA pulldown for PolyG-azidoester modification **(Figures S1-2).**

#### mCXCL10-LPETG

The mature amino acid sequence of murine CXCL10 (aa 22-98) was obtained from NCBI, GeneID 15945. gBlocks were created for this sequence with a C-terminal “LPETG” Sortase recognition site and complementary 5’ and 3’ overhangs to the BamHI/HindIII double digested pCARSF63 Thioredoxin-SUMO fusion expression plasmid (Addgene #64695).^89^ The resulting gBlocks were ligated into pCARSF63 expression plasmids using Gibson assembly (NEB) and transformed into chemically competent SHuffle T7 Express E.coli (NEB). After bacterial liquid cultures were grown to 0.6 OD_600_, protein expression was induced overnight at 16°C with 0.2mM IPTG. The overnight cultures were lysed and sonicated in non-denaturing conditions: B-PER Complete (Thermofisher, 89821), 10 mM imidazole, 0.1% Triton X-100 + COmplete tablet (Millipore Sigma, 11873580001) on ice. mCXCL10-LPETG SUMO fusion proteins were purified by Ni-NTA pulldown and then treated overnight with Endotoxin Removal columns (Thermofisher, 88274). Cleavage of mCXCL10-LPETG from the greater SUMO fusion proteins was carried out via ULP1 digestion (Thermofisher, 12588018) and stored for subsequent PolyG-azidoester modification **(Figures 3A-B, S5)**.

#### Sortase 5M

Bacterial stabs containing Sortase 5M were obtained from Addgene (#51140), deposited by the Ploegh lab.^90^ Individual colonies were picked from plated stabs for 10mL plasmid preps (Invitrogen, K210010), and purified plasmids were transformed into chemically competent T7 Express E. coli (NEB). Cultures were induced with 0.2 mM IPTG overnight at 16 °C, lysed and sonicated on ice in non-denaturing conditions: 20 mM Tris, 125 mM NaCl, 10 mM imidazole, 0.1% Triton X-100 COmplete tablet (Millipore Sigma, 11873580001). Sortase 5M proteins were purified by Ni-NTA pulldown and stored for later use in IgG-LPETG, CCL2 and mCXCL10 sortase tagging experiments.

### Sortase Tagging and Modifying Recombinant Proteins

#### IgG-LPETG (2x)

20 μM IgG-LPETG, 10 μM sortase 5M, and 10mM CaCl^2+^ were mixed with 500 μM of PolyG-4azidoester or 500 μM Triglycine control in sortase reaction buffer (20mM Tris, 50mM NaCl, pH 7.5) for 4 hours at 37°C. Ni-NTA resin was added to remove any unreacted chemokine and sortase 5M. The supernatant, containing pure IgG-4azidoester, was collected, spin-concentrated with 50kDa MWCO columns, and buffer-exchanged into PBS. ^41,46^ Bound IgG was eluted and collected for western blot. Separately, 5μM of purified IgG-LPETG and IgG-4azidoester were mixed with 140 μM DBCO-IR Dye 680 (Licor, 929-50005) at 37°C shaking at 500 RPM overnight. Excess dye was washed away with repeated buffer exchanges into to PBS in 50kDa MWCO spin columns.^41^

#### CCL2-4-azidoester

Ni-NTA resin-bound STEPL fusion proteins containing CCL2-LPETG were incubated with 1mM PolyG-4-azidoester in sortase reaction buffer for 4 hours at 37°C while shaking. Successful sortase tagging releases CCL2-LPETG from the greater fusion protein, leaving the remaining fusion protein bound to Ni-NTA resin. Pure CCL2-4-azidoester was collected in the column flow through, concentrated using 3 kDa molecular weight cut-off (MWCO) columns, and buffer-exchanged into PBS **(Figure S2)**. ^42^

#### mCXCL10-3-azidoester

50 μM of ULP1-digested chemokine was reacted with 10 μM of sortase 5M and 500 μM of PolyG-3-azidoester in sortase reaction buffer for 4 hours at 37°C. Ni-NTA resin was added to remove any unreacted chemokine and sortase 5M. The supernatant, containing pure mCXCL10-3-azidoester, was collected, spin-concentrated with 3kDa MWCO columns, and buffer-exchanged into *in vivo*-grade PBS. ^41^

### Animal and Cell Line Acquisition

#### C57BL/6 mice

C57BL/6 mice were purchased from the Jackson Laboratory (Bar Harbor, ME, USA). All animals were used in accordance with FHCC and SCRI Institutional Animal Care and Use Committee guidelines.

#### B16.F10 melanoma cells

B16.F10 cell lines were obtained from ATCC. These lines were expanded from single clones transduced with lentivirus to express EGF, huCD19, and firefly Luciferase. For tumor assays, 1×10^6^ tumor cells were injected subcutaneously on the right flank of the mouse.

### *In vivo* Hydrogel Studies in C57BL/6 Mice

#### Cortical Hydrogel Injection

A pre-silanized Hamilton Neuros Syringe (Hamilton, 65460-06) was used to aspirate 3µl of chilled 6.5% (w/v) PEG-tetraBCN hydrogel solution, with PEG-Diazide crosslinker added. The solution was promptly injected ∼3mm deep into the parenchyma of the cortex of mice anesthetized with isoflurane. Buprenorphine SR was used as analgesia. After injection, mice were allowed to recover and returned to group housing with no limitations on mobility or access to food and water. Mice were euthanized 7 days after implant and, brains were fixed in 4% neutral-buffered formalin and prepared for IHC.

#### B16 Melanoma Flank Tumor Inoculation and Treatment Administration

C57BL/6 mice were injected subcutaneously with 1 x 10^6^ B16.F10 melanoma cells in 100µL volume in the hind flank. Once tumors were between 200-300 mm^3^ mice were distributed into groups normalized to tumor volume means. Half of the mice were injected intratumorally every other day with murine CXCL10 in PBS (PeproTech, Cranbury, NJ) or a PBS vehicle control. The other half of the mice were injected once with CXCL10-3azidoester-containing hydrogel or an empty hydrogel control. Total injection volumes of 20 µL were used for each treatment group.

### Tissue, Western Blot, and Cell Analysis

#### Locating Injected Gels in Brains via Histology

Mouse brains were harvested, formalin fixed, and paraffin embedded. Brain blocks were then sliced and stained with H&E. Brain sections were imaged using a TISSUEFAX slide scanner (Gnosis) in the imaging core at FHCC.

#### Flow Cytometry Antibodies

All antibodies and viability dyes were purchased from Biolegend (San Diego, CA, USA). Zombie Aqua^TM^ live dead, CD8a (53-6.7), CD4 (RM4-5) and CD3 (17A2). 5 x10^6^ cells were stained for surface or intracellular proteins by incubating cells with antibodies diluted in PBS + 2% BSA for 45 minutes on ice. Cells were then washed 3x in flow cytometry stain buffer and fixed with 2% PFA for 20 minutes prior to acquisition on a LSRII Fortessa (BD Biosciences, San Jose, CA, USA). Samples were analyzed with FlowJo V10 software **(Figure S4)**.

#### Quantifying B16 Tumor Volume and T-cell infiltration

Tumor measurements via caliper began 7 days post-tumor cell injection and were carried out every 2-3 days afterward. Tumor volumes were calculated using the formula (length x width^2^) / 2. At the conclusion of the study, tumors were homogenized into single cell suspensions, stained with a live/dead viability marker and fluorescently conjugated antibodies for CD8, CD4, CD3, and CD44. Cells were analyzed via flow cytometry and total cell counts were calculated by multiplying the target population (i.e., CD8^+^) frequencies of total viable cells by total hemacytometer cell counts. Cell counts were normalized to tumor volume by dividing target populations by tumor volume. **(Figures 5C-D, S5)**

#### Western Blot Analysis

Western blots were run using 4-12% Bis-tris gels (Fisher Sci) in MES running buffer at 180V for 30 mins. Gels were transferred to nitrocellulose using the Iblot or the Iblot 2 semi-dry transfer devices (Fisher Sci). Antibodies and dyes used in this study were: Anti-Human IgG Fc specific (Millipore Sigma I2136-1ML), Anti-6X His tag (Abcam ab9108), Human CCL2/JE/MCP-1 Antibody (RnD Systems, F-279), IRDye 680-DBCO (Licor, 929-50005), Various 680 and 800 channel secondaries (Licor). Blots were scanned and analyzed on the Licor odyssey scanner and software.

### Statistics

Statistics were performed using GraphPad Prism version 9, licensed by SCRI (GraphPad Software, CA, La Jolla, USA). All error bars are specified as either the mean ± standard deviation (SD) or mean ± standard error of the mean (SEM). Hypothesis tests used in this study were: Multiple unpaired t-tests with Holm-Šídák posthoc correction for multiple comparisons, One-Way ANOVA with Tukey’s posthoc correction for multiple comparisons, and Two-Way ANOVA with Holm-Šídák posthoc correction for multiple comparisons. p-Values < 0.05, were considered statistically significant.

## 5. Conclusions

In conclusion, our study demonstrates the successful development of a highly customizable platform for local protein delivery that combines bioorthogonal PEG hydrogels and site-specific protein conjugation via azidoester linkage. Our results show that by modifying the length of the azidoester linker, the release rate of the protein payload can be tailored to suit a range of clinical needs. We also showcase two distinct protein conjugation strategies utilizing the sortase enzyme, Traditional and STEPL, are amenable to a variety of therapeutically relevant proteins and expression systems. Additionally, the sortase and gel functionalization process does not disrupt the biological function of the payload *in vivo*.

The SPAAC-based hydrogel system is injectable, polymerizing rapidly *in situ*, and can be located within tissue days after injection with no noticeable effect on the health of the animal. These features make it particularly well-suited for small, sensitive spaces such as the brain. Furthermore, our study demonstrates one administration of this platform can deliver bioactive murine CXCL10-3azidoester and was successful at recruiting significantly more CD8+ T-cells to a cold melanoma tumor versus controls. Thus, we highlight its potential for clinical applications against human cold tumors.

Overall, our work brings together the fields of chemical engineering, protein engineering, and biology to create a versatile and modifiable platform that can address a range of clinical needs. This platform has the potential to improve current methods of local protein delivery and we believe it will be of great interest to the scientific community working in this field.

## Acknowledgements

Funding for this project was invaluable to our progress and the authors give great thanks to our funding sources: 2017 HHMI Gilliam Fellowship, Alex’s Lemonade Stand, UW/Fred Hutch Interdisciplinary Training Grant T32CA080416, Molecular Medicine Training Program T32GM095421, NIH Diversity Supplement R01CA114567-14, NIH Maximizing Investigators’ Research Award R35GM138036, NIH R01AI132819, NRSA F32CA265056 and NSF DMR 1807398. Many figures in this text were created with Biorender.com, agreement number: WL25ACU57X. Chemical structures were created using ChemDraw ver. 20, licensed by SCRI. The authors attest to no conflicts of interest in this study.

