## Supplemental Data for "Versatile Tissue-Injectable Hydrogels with Extended Hydrolytic Release of Bioactive Protein Therapeutics"

References 15

### General Synthetic Information

Chemical reagents and solvents were purchased from either Sigma-Aldrich or Fisher Scientific and used as received unless otherwise noted. Peptide synthesis reagents were purchased from either ChemPep or Chem-Impex and used as received. Deionized water (d$H_2O$) was generated by a U.S. Filter Corporation Reverse Osmosis System with a Desal membrane. Synthetic chemical reactions were performed under a nitrogen atmosphere in oven-dried glassware and stirred with a Teflon-coated magnetic stir bar unless otherwise noted. Solvents were removed *in vacuo* with a Büchi Rotovapor R-3 equipped with a V-700 vacuum pump and V-855 vacuum controller and a Welch 1400 DuoSeal Belt-Drive high vacuum pump. Microwave-assisted peptide synthesis was performed on a CEM Liberty 1. Semi-preparative reversed-phase high-pressure liquid chromatography (RP-HPLC) was performed on a Dionex Ultimate 3000 equipped with a variable multiple wavelength detector, automated fraction collector, and Thermo 5 µm Synchronis silica 250 x 21.2 mm C18 column. Lyophilization was performed on a LABCONCO FreeZone 2.5 Plus freeze-dryer equipped with a LABCONCO rotary vane 117 vacuum pump. Matrix-assisted laser desorption/ionization time of flight (MALDI-TOF) mass spectrometry was performed in reflectron positive ion mode on a Bruker AutoFlex II using a matrix of α-cyano-4-hydroxycinnamic acid:2,5-dihydroxy benzoic acid (2:1). Fluorescence readings were acquired on a SpectraMax M5 spectrometer using Thermo Scientific Nunc black polypropylene 96-well plates. Protein expression was performed in a Thermo Scientific MaxQ 4000 shaker incubator. Bacterial cells were sonicated using a Qsonica Q500 Sonicator with a 1/4" Microtip Probe.

**Synthesis of Previously Reported Hydrogel Compounds Used in this Work**

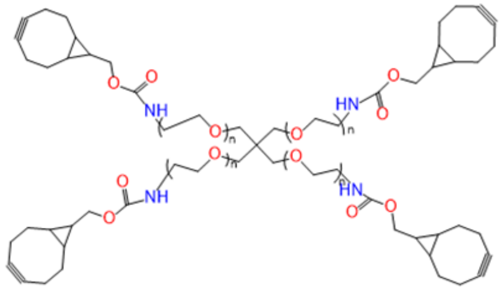

PEG-tetraBCN

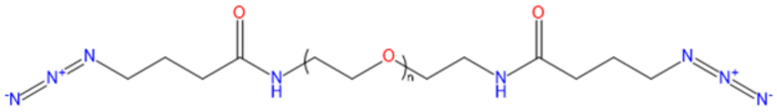

PEG-diazide

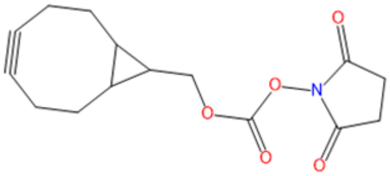

BCN-OSu

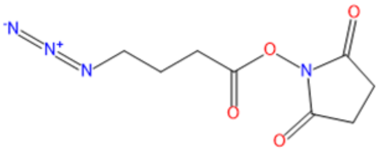

$N_3$-OSu

Poly(ethylene glycol) tetrabicyclononyne (PEG-tetraBCN, $M_n$ ~ 20,000 Da), poly(ethylene glycol) diazide (PEG-diazide, Mn ~ 3,400 Da), (1R,8S,9s)-bicyclo[6.1.0]non-4-yn-9-ylmethyl (2,5-dioxopyrrolidin-1-yl) carbonate (BCN-OSu), and 2,5-dioxopyrrolidin-1-yl 4-azidobutanoate ($N_3$-OSu) were synthesized as previously reported.[1]

**Method S1 Fmoc Solid-Phase Peptide Synthesis**

For the instances when Fmoc-GGGGRS-$NH_2$ was synthesized in-house, a CEM Liberty 1 was used to perform microwave-assisted Fmoc solid phase peptide synthesis (SPPS, 1 mmol scale) to generate the polypeptide. Fmoc deprotection was performed in 20% piperidine (v/v) in dimethylformamide (DMF) with 1-hydroxybenzotriazole (HObT, 0.1 M, 90 °C, 90 sec). Amino acids were coupled to resin-bound peptides upon treatment (75 °C, 5 min) with Fmoc-protected amino acid (2 mmol, 4x), 2-(1H-benzotriazol-1-yl)-1,1,3,3-tetramethyluronium hexafluorophosphate (HBTU, 2 mmol, 4x), and *N*,*N*-diisopropylethylamine (DIEA, 2 mmol, 4x) in a mixture of DMF (9 mL) and *N*-Methyl-2-pyrrolidone (NMP, 2 mL).

### Method S2 Synthesis of Fmoc-GGGGRS-$NH_2$

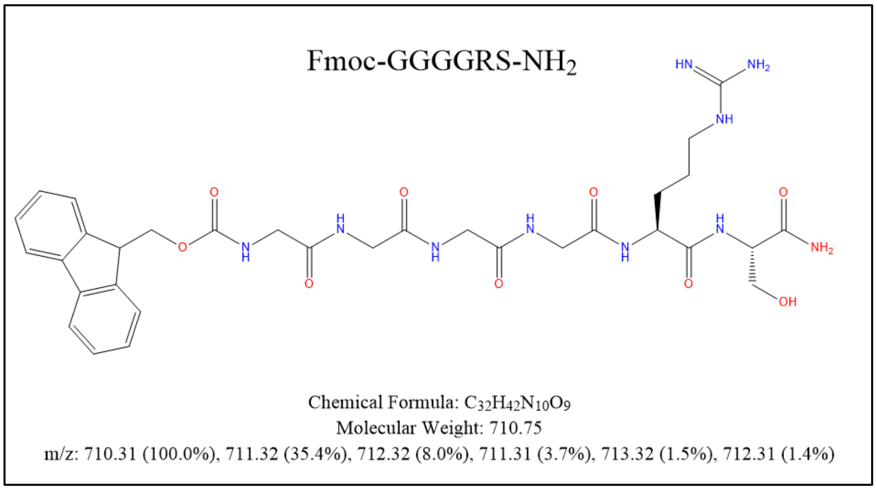

The resin-bound peptide, Fmoc-GGGGRS-NH2, was synthesized by Fmoc SPPS (Method S1) on Rink amide resin (1 mmol scale). Reagents input into the Liberty 1 synthesizer were as follows: 6.24g Fmoc-Arg(pbf)-OH (Chempep, 100202) in 18mL DMF, 5.48g Fmoc-Gly-OH (Chempep, 100801) in 35mL DMF and 1.84g Fmoc-Ser(tbu)-OH (Chempep, 101602) in 9mL DMF. 1.35g of Rink amide at 0.74 scale was used. Deprotection solution: 2.162g HObT dissolved in 160mL of 20% Piperidine in DMF. Activator Base: 10.5mL DIEA in 20mL NMP. Activator: 9.48g HBTU dissolved in 50mL DMF. After the synthesis program was completed, the resin was rinsed 3x in DCM. The peptide was cleaved from the resin using 40mL Cleavage Cocktail (95% Trifluoroacetic Acid, 5% $H_2O$, 5% Triisopropylsilane) stirring for 2 hours at RT. This cleavage solution was crashed in Di-ethyl Ether (2x), spun down and dried under a nitrogen atmosphere. The crude peptide was purified via RP-HPLC using a 55-minute gradient from 20-100% acetonitrile:H2O; lyophilization yielded the final product (Fmoc-GGGGRS-NH2) as a white solid. Peptide purity was confirmed using MALDI-TOF. Expected MW: 710.31 Da. Actual: 711.30 Da

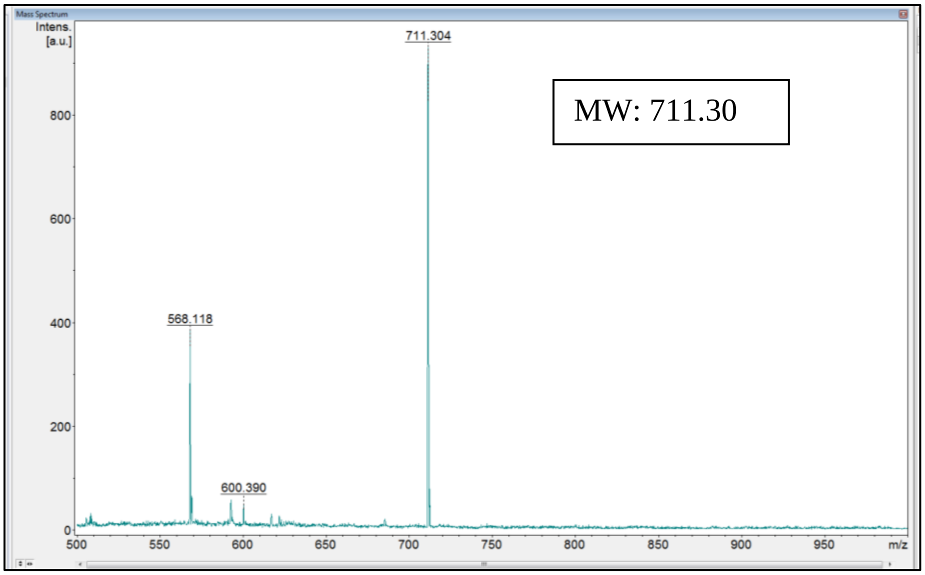

**Method S3 Synthesis of DEAC-OH**

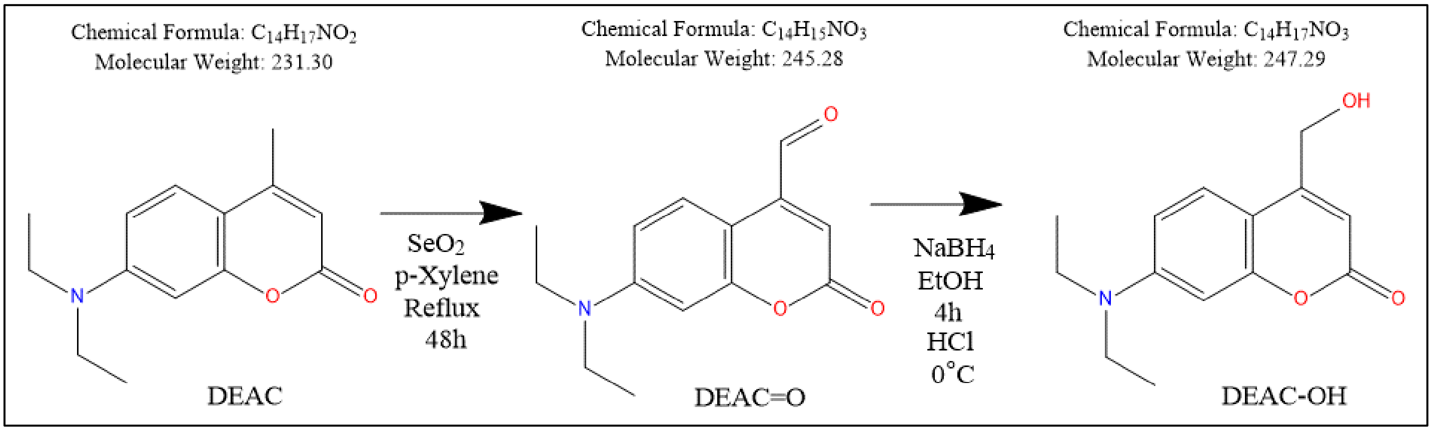

Simplified synthesis scheme of 7-Diethylamino-4-(hydroxymethyl) coumarin (DEAC-OH, MW: 247.29 Da) previously synthesized in-house utilizing 7-Diethylamino-4-methylcoumarin (DEAC, 231.29 Da) (Fisher Sci, 50534181) as starting material. Briefly, DEAC (0.5 g; 2.16 mmol) was combined with selenium dioxide (0.7 g; 6.3 mmol; 3 eq) in a round bottom flask and dissolved in p-Xylene (30 mL). The solution was refluxed for 48 hours under vigorous stirring and quickly became dark brown over the course of 3 hours. After 48 hours, the mixture was filtered and concentrated under reduced pressure. The residual brown oil was dissolved in Ethanol and Sodium Borohydride (0.985 g; 26 mmol) was added. The solution was stirred for 4 hours before careful hydrolysis with HCl (1 M; 2.5 mL) at 0 °C. The solution was then diluted into water and extracted with 3 portions of dichloromethane before drying with $MgSO_4$, filtration, and concentration under reduced pressure. The result was purified using a 0-70% gradient of ethyl acetate in hexanes to provide the product, DEAC-OH (290 mg; 54% yield).

### Figure S1 Amino Acid Sequence of Human CCL2 STEPL Fusion Protein

*hCCL2*-*LPETG*-*Sa-SrtA*-*6xHis*:

QPDAINAPVTCCYNFTNRKISVQRLASYRRITSSKCPKEAVIFKTIVAKEICADPKQKWVQDSMDHLDKQTQTPKTL
ELPETGGGSGGSGGSGGSGGSQAKPQIPKDKSKVAGYIEIPDADIKEPVYPGPATPEQLNRGVSFAEENESLDDQNI
SIAGHTFIDRPNYQFTNLKAAKKGSMVYFKVGNETRKYKMTSIRDVKPTDVEVLDEQKGKDKQLTLITCDDYNEKTG
VWEKRKIFVATEVKHHHHHH

The complete amino acid sequence of mature human CCL2 (aa 24-99) fused with sortase in the pSTEPL expression plasmid.[2,3] The sequence begins with CCL2 in red, separated from the rest of the fusion protein by a $GGS_5$ linker (black). The LPETG sortase recognition motif is labeled in blue, a truncated sortase enzyme is labeled in green, and a C-terminal 6x-His tag is labeled in orange. It should be noted that bacterial methionine aminopeptidase (MAP) may not always remove the formyl-methionine used to initiate translation of this protein.[4] Although this is not a problem for every protein, chemokine-receptor signaling is dependent upon the preservation of the native N-terminus (i.e.: H-Gln).[5] We did not perform receptor-mediated assays with CCL2, but addressing this was necessary for our studies utilizing CXCL10 in **Figure S4**.

**Figure S2 Conjugating PolyG-4azidoester to Human CCL2 via STEPL**

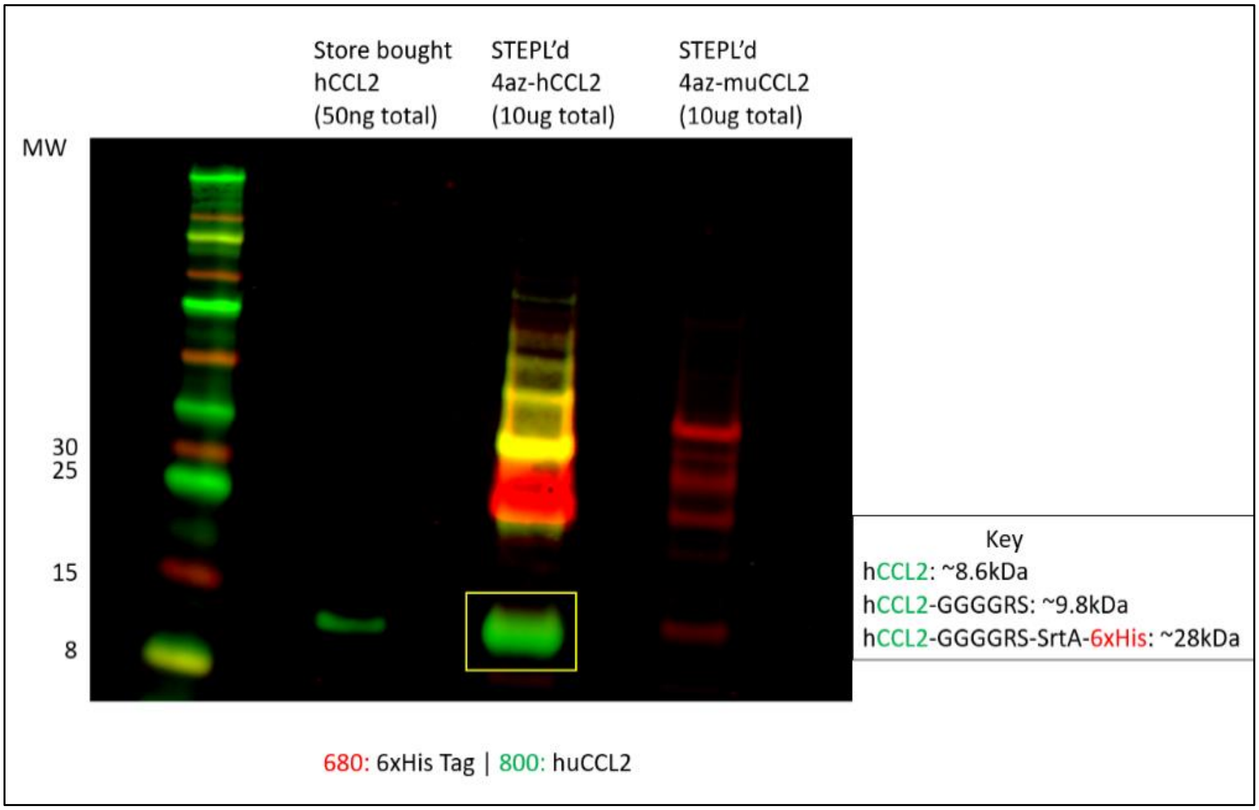

A Western blot was performed to confirm the expression and modification of the human CCL2 pSTEPL fusion protein with PolyG-4azidoester attachments. The blot consisted of three lanes: a positive control (store-bought human CCL2, Thermofisher RP-8648), a collected flow-through from an on-column reaction between human CCL2-STEPL with PolyG-4azidoester, and a collected flow-through from an on-column reaction between murine CCL2-STEPL with PolyG-4azidoester as a negative control. The left lane showed the positive control detected by a polyclonal human CCL2 antibody (RnD Systems, AF-279-SP) in the green channel. The middle column showed the human CCL2 separated from the greater STEPL fusion protein and visible in the green channel (highlighted in a yellow box). The right column showed the negative control, with murine CCL2 labeled non-specifically with the 6x-His Tag antibody (Abcam, ab18184) in the red channel but not detected by the human-specific CCL2 antibody.

**Figure S3 MALDI-TOF Confirmation of PolyG-3,4azidoester Synthesis**

<u>**Expected**</u> <u>**Actual**</u>

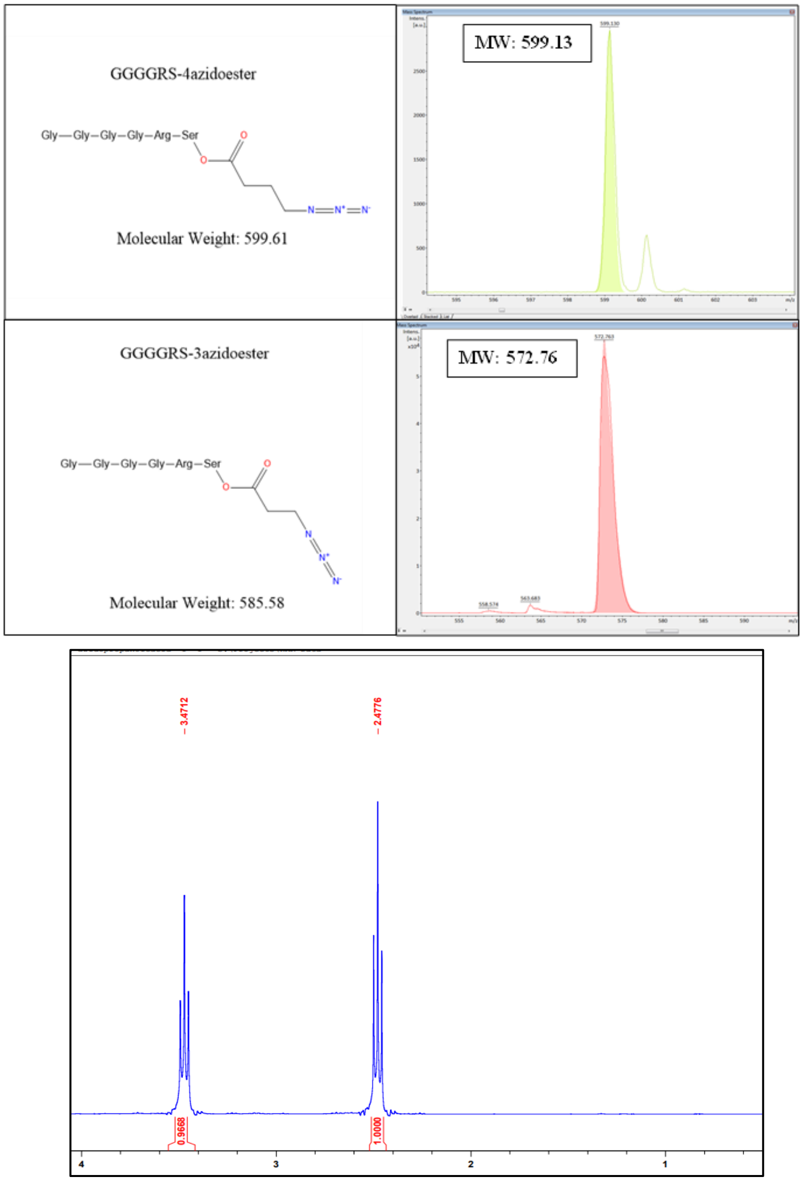

MALDI-TOF mass spectrum traces were obtained for each of the synthesized products. PolyG-4azidoester was found to match its expected molecular weight (MW: 599.13 Da), and the resulting lyophilized product appeared as a light-yellow oil, as anticipated. However, PolyG-3azidoester appeared to be missing approximately 14 Da, making its molecular weight closer to that of PolyG-2azidoester (MW: 571.56 Da) rather than its own expected molecular weight (MW: 585.58 Da). H-NMR on our stock of 3-azidopropanic demonstrates the presence of two triplets at 3.47ppm and 2.47ppm, confirming the identity of 3-azidopropanic acid. We conclude the observed mass discrepancy may be an artifact of the spectroscopy process.

### Figure S4 Amino Acid Sequence, Purification and Sortag of mCXCL10 SUMO Fusion

*6xHis-TrxA- SUMO-mCXCL10-LPETG-Strep*:

HHHHHHGSDKIIHLTDDSFDTDVLKADGAILVDFWAEWCGPCKMIAPILDEIADEYQGKLTVAKLNIDQNPGTAPKY
GIRGIPTLLLFKNGEVAATKVGALSKGQLKEFLDANLAGTSDSEVNQEAKPEVKPEVKPETHINLKVSDGSSEIFFK
IKKTTPLRRLMEAFAKRQGKEMDSLRFLYDGIRIQADQTPEDLDMEDNDIIETHREQIGGIPLARTVRCNCIHIDDG
PVRMRAIGKLEIIPASLSCPRVEIIATMKKNDEQRCLNPESKTIKNLMKAFSQKRSKRAPGGSGGSLPETGWSHPQF
EK

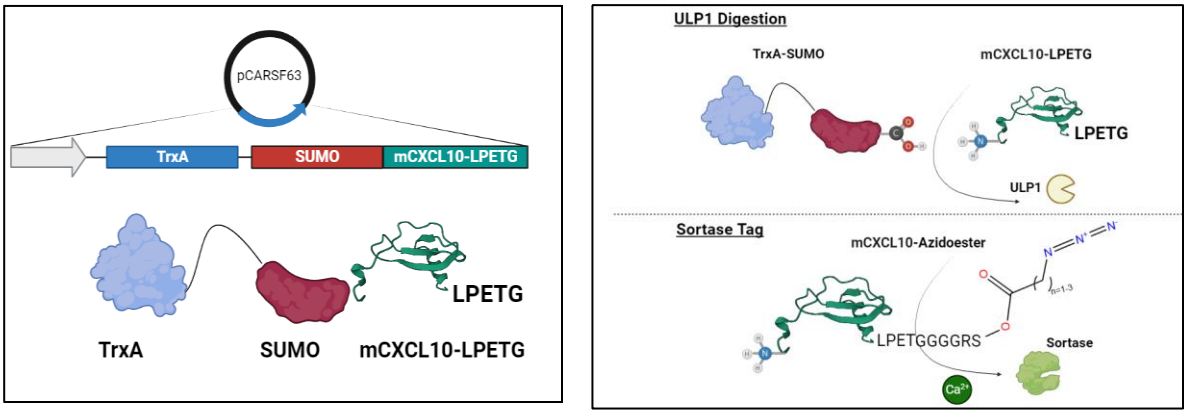

The complete amino acid sequence of mature, murine CXCL10 (aa 22-98) fused with Thioredoxin and SUMO within the pCARSF63 expression plasmid is shown.[6] The fusion protein begins with an N-terminal 6x-His tag in orange, followed by Thioredoxin in purple, and then SUMO in Green. The terminal GG of SUMO are subject to specific cleavage by SUMO protease (ULP1), releasing what immediately follows with a native N-terminus (i.e., H-Ile). The sequence of murine CXCL10 is labeled in red, followed by LPETG in light blue and a Strep Tag in navy blue. Upon overnight digestion of this fusion protein by ULP1, the resulting chemokine has a native N-terminus which preserves receptor signaling and biological activity. We then performed traditional sortase tagging to add a PolyG-azidoester to its C-terminus.

**Figure S5 SDS-PAGE Confirmation of Purified mCXCL10-TrxA SUMO Fusion Protein**

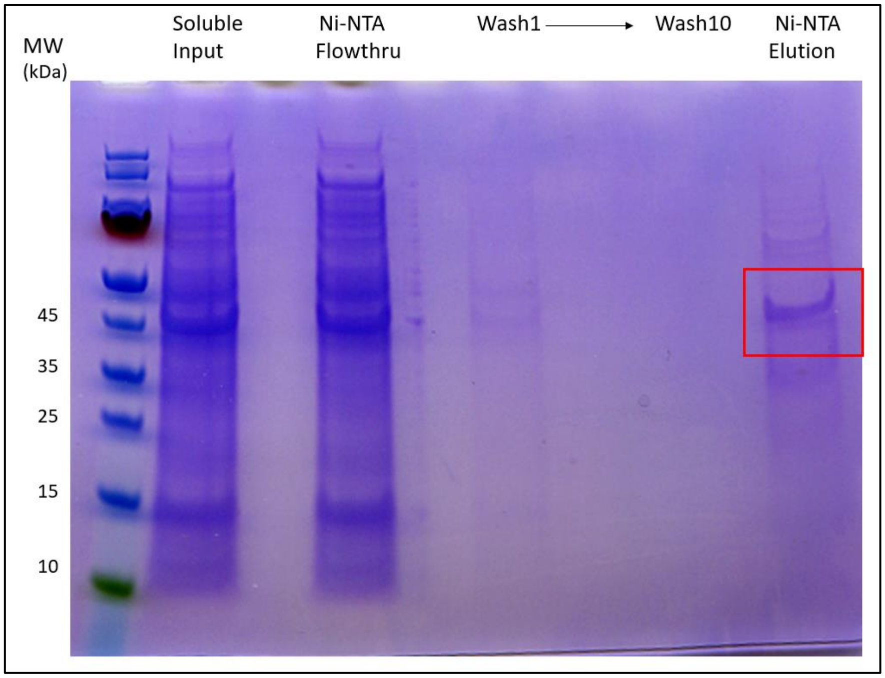

This SDS-PAGE gel depicts the purification process after the expression of the SUMO fusion protein in Shuffle T7 Express *E. coli*. The purification involves a Ni-NTA pulldown and repeated wash steps. The purified fusion protein is collected in the Ni-NTA elution lane on the far right, which is highlighted in a red box. The purified protein will then be stored for future ULP1 cleavage and sortagging.

**Figure S6 Flow Cytometry Gating Scheme for CD4$^{+}$ and CD8$^{+}$ T-cells**

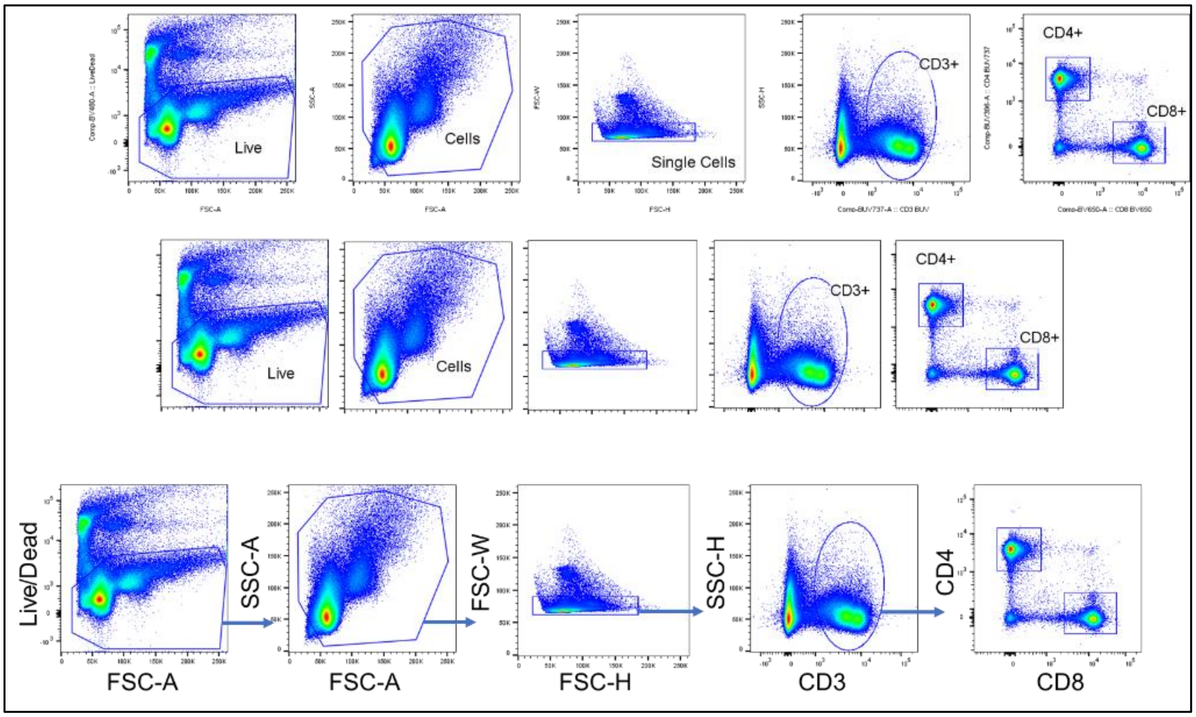

The image depicts the gating scheme used to identify T-cell populations in B16 melanoma flank tumors at the conclusion of our *in vivo* study. All antibodies and viability dyes were purchased from Biolegend: Zombie Aqua$^{TM}$ live dead, CD8a (53-6.7), CD4 (RM4-5), and CD3 (17A2). 5 x$10^{6}$ cells were stained for surface or intracellular proteins by incubating cells with antibodies diluted in PBS + 2% BSA for 45 minutes on ice. Cells were then washed 3x in flow cytometry stain buffer and fixed with 2% PFA for 20 minutes prior to acquisition on a LSRII Fortessa (BD Biosciences). Samples were analyzed with FlowJo V10 software.

**Figure S7 Quantifying CD4+ Infiltration into Flank Tumors**

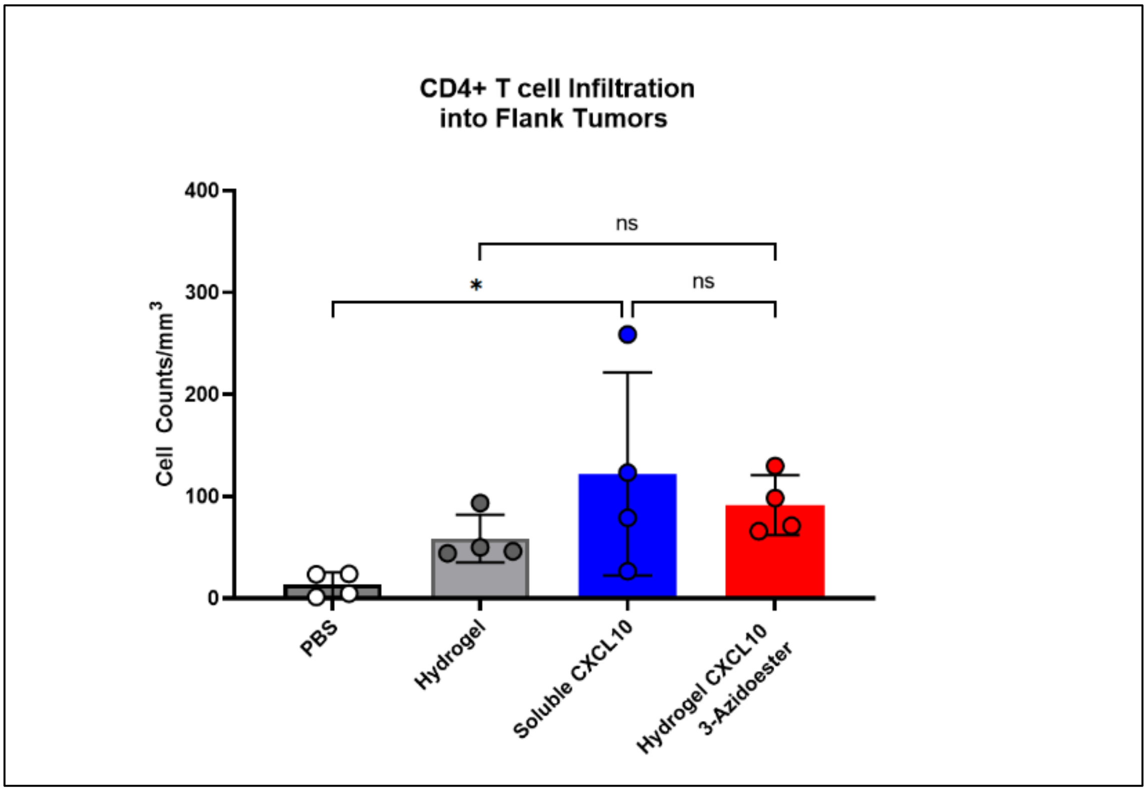

The data presented in this graph indicates that the hydrogel CXCL10-3-azidoester treatment does not result in a significant difference in $CD4^+$ infiltrate compared to the empty hydrogel alone. However, CXCL10 solutions in PBS did show a significant difference when compared to PBS alone. There was no significant difference in $CD4^+$ cells between CXCL10 in PBS and CXCL10-3azidoester hydrogels. Data are presented as mean ± SD for n=4 replicates. Statistical significance was determined by ordinary one-way ANOVA followed by Holm-Šídák posthoc correction. [(ns) not significant, (*) $p < 0.05$].
